# The oral pathogen *Porphyromonas gingivalis* resists the antimicrobial peptide LGL13K and evades the D-enantiomer by synonymous mutations in *hagA*

**DOI:** 10.1101/2023.09.07.556717

**Authors:** Sven-Ulrik Gorr, Ruoqiong Chen, Juan E. Abrahante, Paul B.M. Joyce

## Abstract

*Porphyromonas gingivalis* is a keystone pathogen for periodontal disease. The bacteria are black-pigmented and require heme for growth. *P. gingivalis* exhibit resistance to many antimicrobial peptides, including the L-enantiomer of the antimicrobial peptide GL13K, which contributes to their success in the oral cavity. *P. gingivalis* W50 was resistant to LGL13K but susceptible to the stereo-isomer DGL13K. Upon prolonged exposure to DGL13K, a novel non-pigmented mutant was isolated that showed a low minimum inhibitory concentration and two-fold extended minimum duration for killing by DGL13K, consistent with tolerance to this peptide. The DGL13K tolerant bacteria exhibited synonymous mutations in the *hagA* gene. The mutations did not prevent mRNA expression but were predicted to alter mRNA structure. The non-pigmented bacteria were deficient in hemagglutination and hemoglobin binding, suggesting that the HagA protein was not expressed. This was supported by whole cell ELISA and gingipain activity assays, which suggested the absence of HagA but not two closely related gingipains. *In vivo* virulence was similar for wild-type and non-pigmented bacteria in the *Galleria mellonella* model. Loss of the hemagglutinin HagA may allow bacteria to escape from a biofilm that is under attack by antimicrobial peptides.

## Introduction

Periodontitis is one of the most common infectious diseases in middle-aged and older adults, with a prevalence of 50% in the U.S. (1). Periodontitis has consistently been associated with a number of oral bacteria, including *Porphyromonas gingivalis* (2–4), which is considered a keystone pathogen that can initiate dysbiosis while remaining at relatively low frequency in the healthy oral microbiome (2, 4, 5). *P. gingivalis* infections have been linked to several systemic diseases (6), including atherosclerosis (7, 8), rheumatoid arthritis (9), and Alzheimer’s disease (7, 10), although causation is still debated (11–14).

*P. gingivalis* are black-pigmented bacteria that require heme for growth (15–17). The bacteria produce multiple virulence factors that affect the host inflammatory response and contribute to the destruction of periodontal tissues supporting the teeth (18). Among these virulence factors are three cysteine-proteases, Lys-gingipain (Kgp), Arg-gingipain A and B (RgpA and RgpB) (19) and a related hemagglutinin A (HagA) (18). The latter shares adhesion domains with the gingipains Kgp and RgpA but lacks proteolytic activity (20). These proteins form a cell-surface complex (21) that is associated with heme-acquisition and colony pigmentation (16, 17).

A large number of non-pigmented mutants of *P. gingivalis* have been described (16, 22–31). The mutations typically affect the gingipain-hemagglutinin complex (23, 26, 28, 30–32), other type IX secretion system (T9SS) components (22, 33), or LPS structure (24, 25, 27). As a result, non-pigmented mutants may be less virulent than WT bacteria (29).

*P. gingivalis* thrives in the oral cavity, an environment that is rich in antimicrobial peptides (AMPs), both above and below the gum-line (34, 35). The success of *P. gingivalis* in this environment can be attributed to the formation of biofilms (36), invasion of epithelial cells (37–39) and resistance to AMPs (34, 40–45). The latter is likely mediated by modification of the lipopolysaccharide (LPS) structure in the outer membrane (40, 46, 47), which can generate a less negatively charged surface that does not attract cationic peptides (40). In addition, gingipains can contribute to resistance (42, 46), although this may be peptide specific (41).

We have developed the AMP GL13K (43, 48, 49), which is derived from the sequence of the salivary protein BPIFA2 (BPI fold-containing family A member 2; former names: parotid secretory protein, PSP, SPLUNC2, C20orf70) (50–52). The L-amino acid version of this peptide, LGL13K, shows activity against the Gram-negative bacteria *Escherichia coli* and *Pseudomonas aeruginosa* (43) and biofilms of the latter (53). LGL13K is degraded by bacterial proteases (53, 54) and the peptide is not active against *P. gingivalis* strains 55977, W50, and DPG3 (43). To explore further the relative resistance of *P. gingivalis* to this peptide, we compared the activity of LGL13K to that of a protease-resistant D-enantiomer, DGL13K (53, 54). The results suggest that, unlike LGL13K, DGL13K can defeat multiple bacterial resistance mechanisms but bacteria can gain tolerance to DGL13K through mutations in the *hagA* gene.

## Results

We have previously reported that three strains of *P. gingivalis* (W50, ATCC 53977 and DPG3) are resistant to LGL13K but not to the endogenous antimicrobial peptide LL-37 (43). We have since designed an all-D amino acid version of this peptide (DGL13K) (53), which resists proteolytic degradation and kills both Gram negative and Gram positive bacteria (53–56). Timed-kill assays showed that *P. gingivalis* W50 were killed by DGL13K but not LGL13K or polymyxin B (**Fig. 1A**). In the course of DGL13K-treatment, a non-pigmented variant of W50 emerged and was dominant prior to complete killing of the bacteria (**Fig. 1A**). Thus, the minimum duration for killing (MDK) (57) was about 60 min for pigmented colonies and 120 min for non-pigmented colonies (**Fig. 1B**).

**Figure 1.**
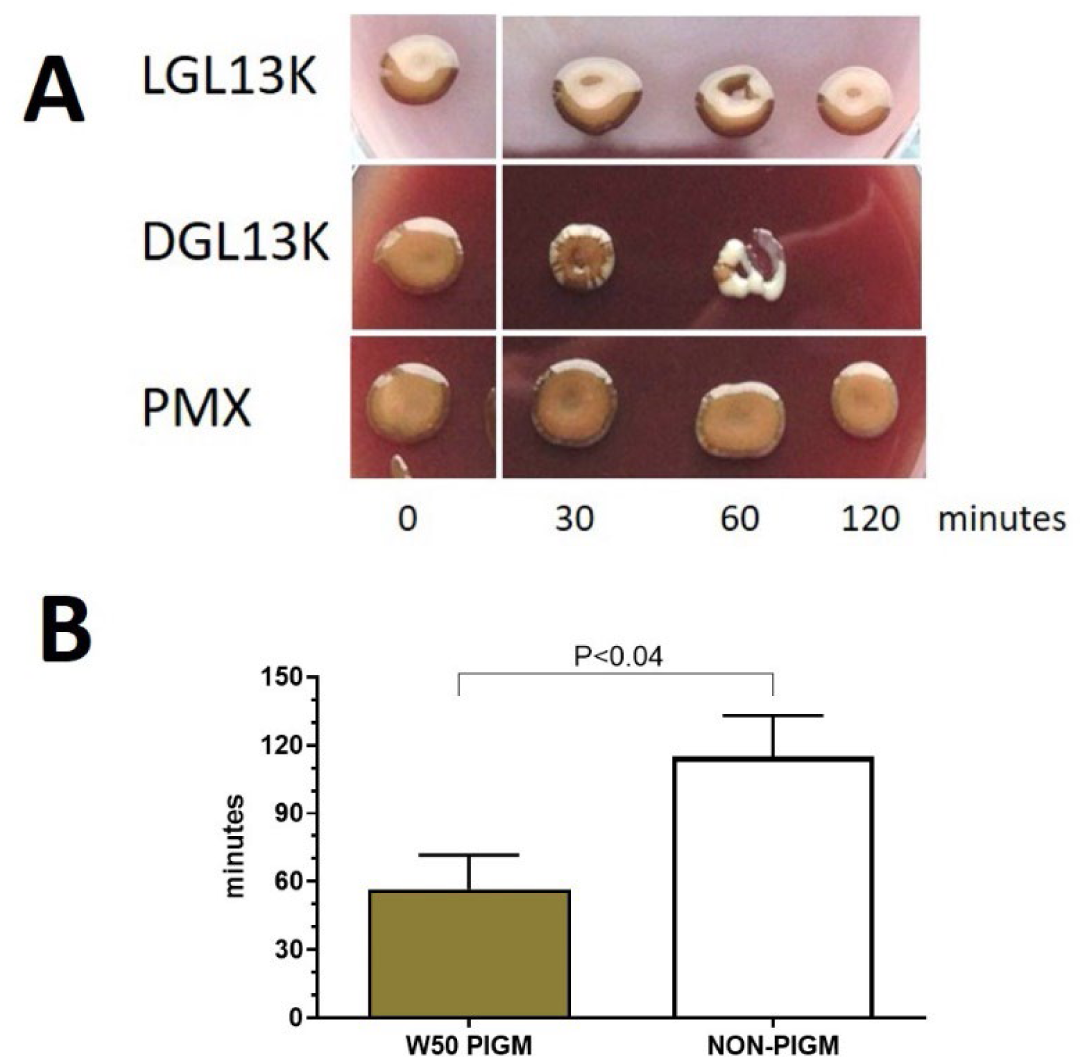
Time course of bacterial killing in solution. **A**. *P. gingivalis* W50 were incubated for 2 hours with LGL13K, DGL13K or polymyxin B (PMX) (100 µg/ml) and aliquots plated on blood agar at the times indicated. The figure is a composite of separate culture pates from a single representative experiment. **B**. Minimum duration for killing (MDK) was calculated for pigmented W50 and non-pigmented W50 (Non-pigm). Data from five independent experiments are shown as mean ± SEM. N=6. The two groups were compared by unpaired Student’s t-test.

Non-pigmented colonies from the DGL13K selection experiments, were further expanded in culture medium, in the absence of peptide. The minimum inhibitory concentrations (MICs) were determined for WT *P. gingivalis* and the non-pigmented W50 isolate. LGL13K (up to 1024 µg/ml) did not inhibit growth of any of the tested strains (**Table 1**), consistent with the timed-kill assays shown above. In contrast, DGL13K similarly inhibited the growth of WT and the non-pigmented isolate of W50.

**Table 1.** Minimal Inhibitory Concentrations of DGL13K, LGL13K and polymyxin B against WT *P. gingivalis* strain W50 and the Non-pigmented W50 isolate. MICs were expressed as mean ± SEM (N). >1024 – growth was not inhibited at the highest peptide concentration tested. The data are from 5-6 independent experiments, each with 2-6 replicates.

| Peptide | DGL13K | LGL13K | Polymyxin B |
| --- | --- | --- | --- |
| Strain | Mean MIC ± SEM (N) | | |
| WT | 11 ± 2 μg/ml (17) | >1024 μg/ml (18) | 74 ± 13 μg/ml (10) |
| Non-pigmented | 11 ± 1 μg/ml (15) | >1024 μg/ml (12) | 61 ± 13 μg/ml (10) |

### Genome sequence and transcription

To investigate the molecular mechanism behind the lack of colony pigmentation and increased MDK for bacteria treated with DGL13K, non-pigmented colonies from four independent experiments were expanded and analyzed by whole-genome sequencing. The untreated W50 WT strain differed from the W83 reference genome sequence (NCBI Reference Sequence: NC_002950.2) in only three locations (61429, 1802991 and 1803013; W83 numbering system).

In addition, sequence analyses revealed seven mutations that were present in at least 75% of sequence reads from DGL13K-treated samples but not in WT controls. Two of these mutations were found in two of four experiments (**Table 2**) and were selected for further analysis.

**Table 2.** SNPs in DGL13K-treated non-pigmented *P. gingivalis* W50 (W50 *hagA23/167*).

| **Genome location** | **Gene** | **Codon polymorphism**<br>**pigmented → non-pigmented** | **Protein sequence** |
| --- | --- | --- | --- |
| 1936023 | *hagA* | TCG → TCA | Ser → Ser |
| 1936167 | *hagA* | TAT → TAC | Tyr → Tyr |

Both were synonymous SNPs located in the *hagA* gene. The corresponding non-pigmented mutant was named W50 *hagA23/167* to denote the locations of the two SNPs. The two mutations did not prevent *hagA* transcription since the corresponding mRNA was readily identified by RT-PCR with no consistent differences between WT and non-pigmented bacteria (**Fig. 2**).

**Figure 2.**
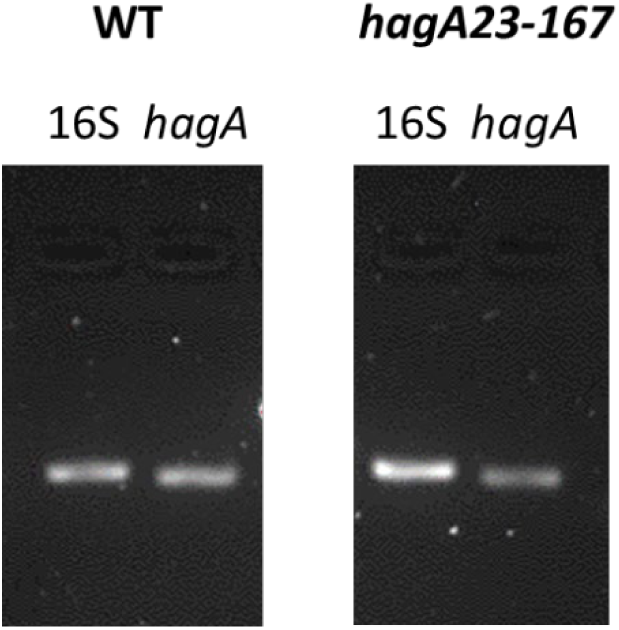
RT-PCR of *P. gingivalis* W50 WT and non-pigmented isolate after DGL13K treatment (*hagA23/167*). Transcripts of 16S rRNA and *hagA* were amplified by RT-PCR and visualized by agarose gel electrophoresis. The images are representative of four (WT) and six (*hagA23/167*) independent samples.

Codon usage can affect both transcription and translation (58). Analysis of codon usage in *hagA* revealed that all six Ser codons and both Tyr codons were used. A T→C transition in the *hagA23/167* sequence changed the more frequently used TAT Tyr codon (67 of the 100 Tyr codons in the WT sequence) to the less frequently used TAC Tyr codon (33 codons in the WT sequence) in *hagA23/167*. The G→A transition in the *hagA23/167* sequence converted the low-use TCG Ser codon (8 of the 131 Ser codons in the WT sequence) to the 3-times more frequently used TCA Ser codon (23 codons in the WT sequence) (**Table 3**).

**Table 3.** Codon usage for Ser and Tyr residues in *hagA*. The number of uses for each codon in WT *hagA* are shown. The WT (shaded) and mutated codons (bold italic) are marked for each SNP.

| Codons | Ser | Tyr |
| --- | --- | --- |
| TCT | 52 |  |
| TCC | 28 |  |
| **TCA** | **23** |  |
| TCG | 8 |  |
| AGC | 7 |  |
| AGT | 13 |  |
| TAT |  | 67 |
| **TAC** |  | **33** |
| Total | 131 | 100 |

### mRNA structure prediction

In addition to codon usage, mRNA structure can affect translation efficiency (59). To determine if the synonymous SNPs affected predicted mRNA structure, the WT and *hagA23/167* sequences were modeled in a 493 nucleotide domain representing the C-terminal cleaved adhesion domain (orphan K3 domain (60)), which contains both synonymous SNPs. The predicted structure of the *hagA23/167* mRNA differed substantially from the WT mRNA (**Fig 3 A,D**). Hypothetical mRNA structures, which contained only one of the two synonymous SNPs were modeled (**Fig. 3B-C**) to determine if both SNPs contributed to the predicted structure of *hagA23/167* mRNA. Based on this analysis, it appears that alteration of the Ser codon at position 1936023 (**Fig. 3C**) causes most of the structural change determined in the *hagA23/167* mRNA. The same conclusion was reached when the secondary structures were modeled by predicting the minimum free energy (MFE) (data not shown), which strengthens the reliability of the predicted structures (61).

**Figure 3.**
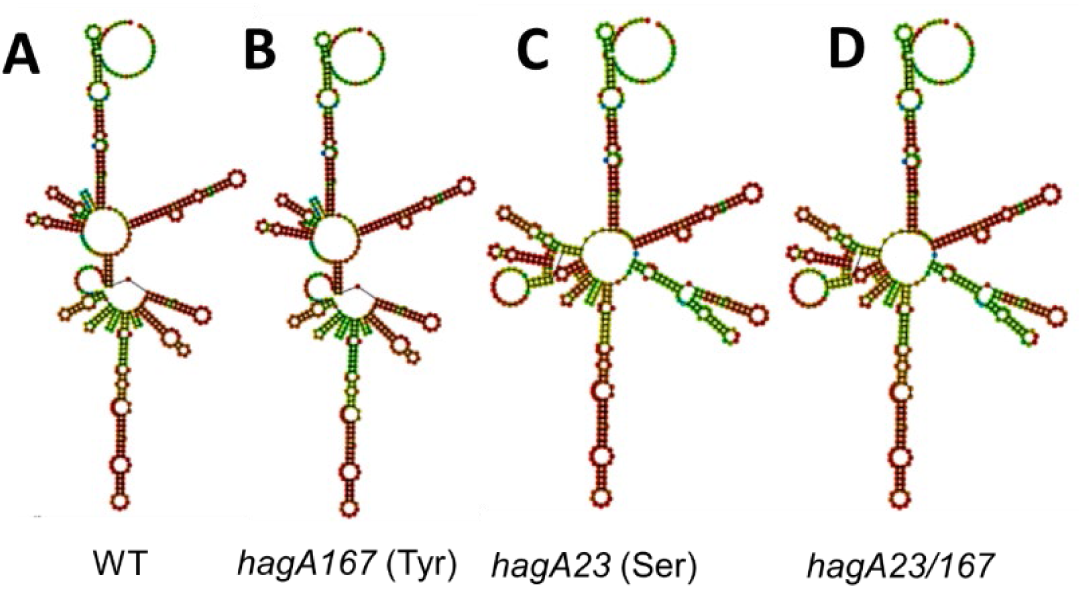
Predicted structure of a 493 nucleotide segment of the mRNA sequence coding for the C-terminal K3 cleaved adhesin domain of *hagA.* **A.** WT; **B.** Model of a hypothetical *hagA167* mutant, which was altered at the Tyr codon at position 1936167. **C**. Model of a hypothetical *hagA23* mutant, which was altered at the Ser codon at position 1936023. **D.** Model of a hypothetical *hagA23/167* double mutant.

### HagA expression and function

Heme-binding and hemagglutination are associated with proteins that contain the Cleaved Adhesin Domain (CAD) (20), including Lys-gingipain (Kgp), Arg-gingipain A (RgpA) and hemagglutinin A (HagA). The non-pigmented phenotype of the non-pigmented *hagA23/167* mutant suggested that it could be deficient in heme binding and uptake, which was confirmed experimentally (**Fig. 4**). Thus, the *hagA23/167* mutant showed deficient heme-binding (**Fig. 4A**) and hemagglutination (**Fig. 4B**), consistent with the non-pigmented phenotype.

**Figure 4.**
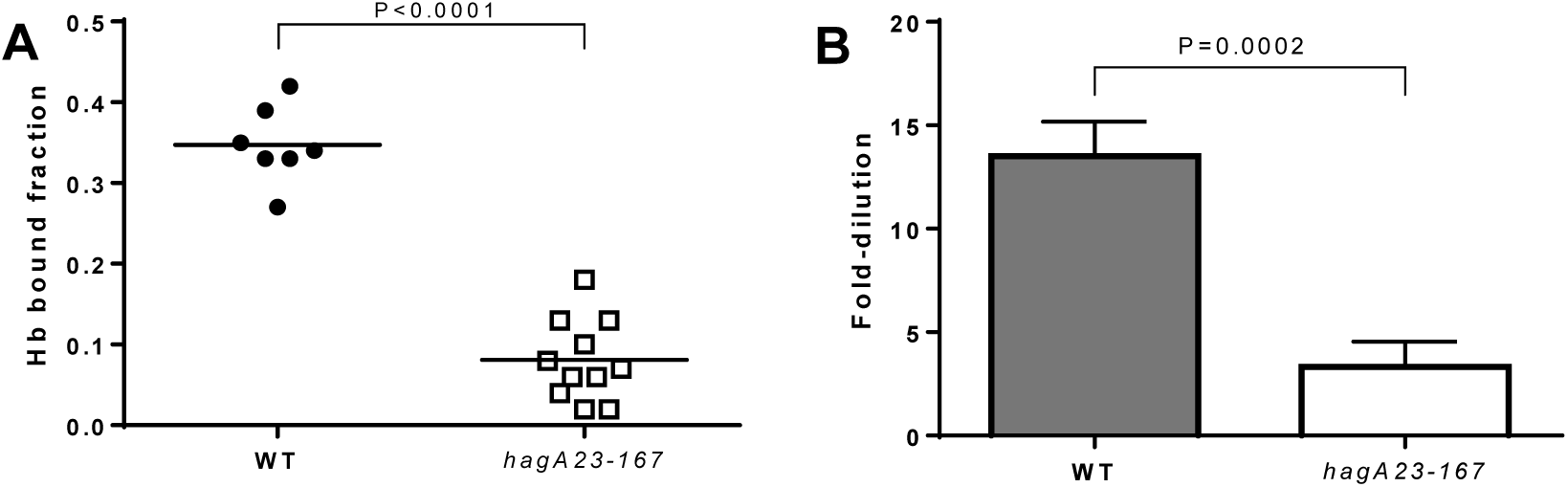
**A**. Hemoglobin-binding of *P gingivalis*. W50 (WT) and non-pigmented W50 (*hagA23/167*) cells were incubated for 60 min with hemoglobin and the bound fraction determined. Data from three independent experiments were analyzed by Student’s t-test, N=7-11. Line indicates group mean. **B**. <u>Hemagglutination by *P. gingivalis*</u>. Two-fold serial dilutions of W50 (WT) and non-pigmented W50 (*hagA23/167*) were incubated with red blood cells and hemagglutination determined by the matting of red blood cells. The maximal bacterial dilution that resulted in hemagglutination was determined in five independent experiments and analyzed by Student’s t-test. Data are shown as mean ± SEM, N=8-9.

To determine if the Cleaved Adhesin Domain proteins were expressed at the cell surface, the monoclonal antibody 61BG1.3, which recognizes a hemagglutinating epitope in these proteins (20), was used for whole-cell ELISA (**Fig. 5**). WT *P. gingivalis* resulted in a strong antibody signal, consistent with the expression of all three proteins containing this epitope, while the *hagA23/167* mutant showed reduced binding, consistent with the absence of one or more CAD proteins.

**Figure 5.**
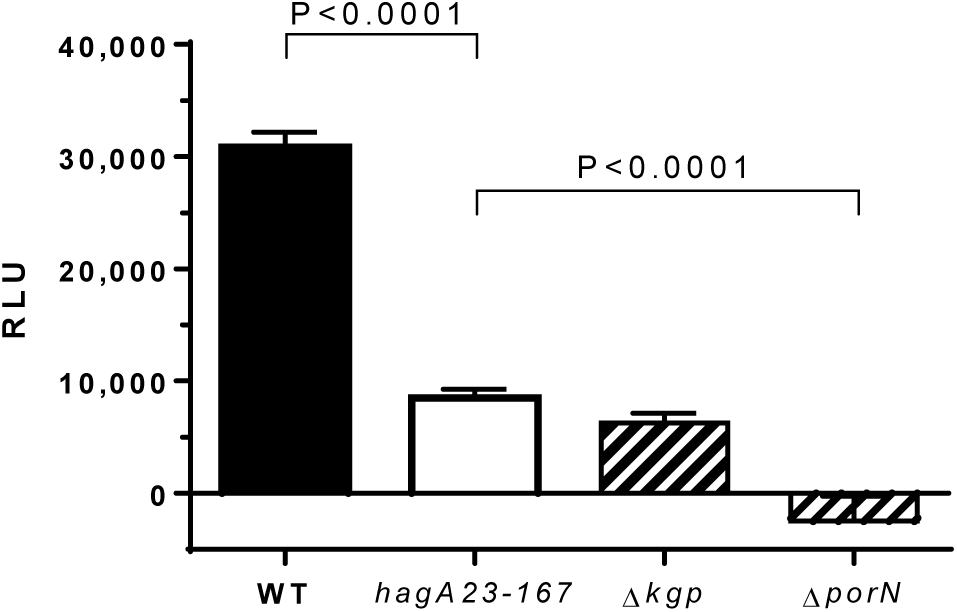
Whole cell ELISA of *P. gingivalis*. Biofilms of *P. gingivalis* W50 (WT), *hagA23/167*, Δ*kgp* or Δ*porN* were cultured on hydroxyapatite-coated pegs. The surface expression of Kgp, RgpA and HagA was determined by immunolabeling and quantitated by chemiluminescence (RLU). Statistical outliers were removed and the data from two independent experiments were expressed as mean ± SEM. The data for *hagA23/167* were compared to control groups by one-way ANOVA with Dunnett’s multiple comparison post-test, N=16.

For comparison, the binding of 61BG1.3 was also tested with two mutants of the closely related *P. gingivalis* strain W83. As expected (62), the secretion mutant Δ*porN,* did not express the three proteins at the cell surface. The mutant Δ*kgp*, which does not express the Lys-gingipain, exhibited intermediate antigen expression levels, similar to that observed for *hagA23/167* mutant (**Fig. 5**). These data suggest that the *hagA23/167* mutant lacks surface expression of one of the CAD proteins.

### Gingipain activity

To distinguish between HagA and Lys-gingipain expression, we tested if the mutant cells possessed hemoglobinase activity, a measure of the Lys-gingipain Kgp (23). **Fig 6** shows that the hemoglobinase activity in the *hagA23/167* mutant did not differ from WT cells, suggesting that the latter mutation does not affect Lys-gingipain expression. This interpretation was confirmed with the Lys-gingipain deficient strain Δ*kgp*, which showed a significant reduction of hemoglobinase activity (**Fig. 6**).

**Figure 6.**
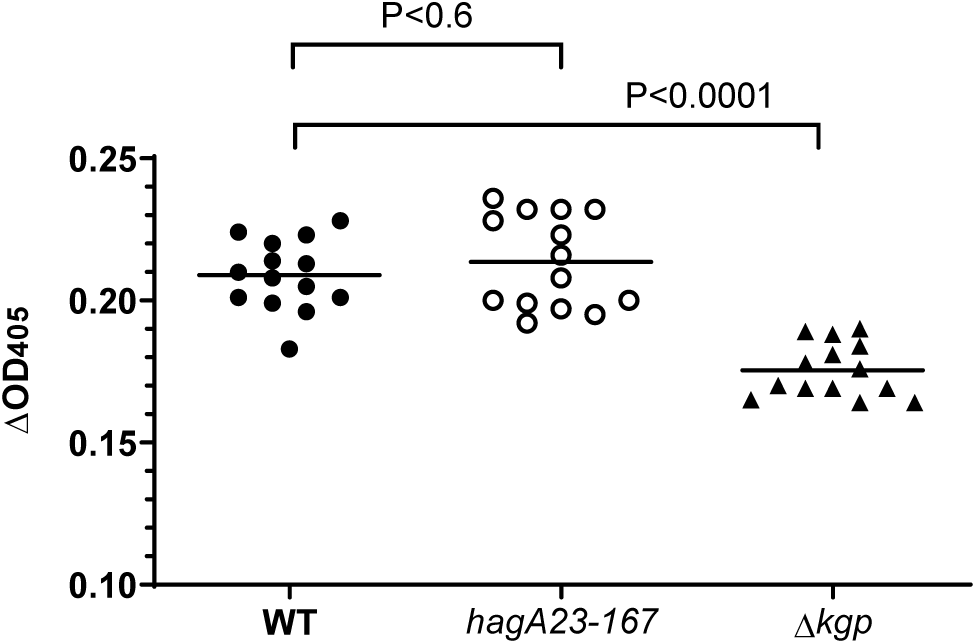
Hemoglobinase activity in *P. gingivalis*. Secretion medium from WT, the *hagA23/167* mutant or Lys-gingipain mutant (Δ*kgp*) were incubated with hemoglobin and the decrease in OD_405_ recorded after 24h, as a measure of hemoglobinase activity. Data from two independent experiments were adjusted for differences in initial culture density between strains and analyzed by one-way ANOVA with Dunnett’s post-test. N=14. Lines indicate the mean of each group.

### In vivo virulence

The above results suggested that *hagA23/167* does not express HagA at the cell surface. To test if depletion of this virulence factor (63) affected bacterial virulence, WT and mutant strains were injected into *Galleria mellonella*, an in vivo model of bacterial infection (64, 65). The in vivo virulence of *hagA23/167* is similar to that of WT W50 (**Fig 7**). Both strains show significant virulence compared to larvae that were injected with PBS.

**Figure 7.**
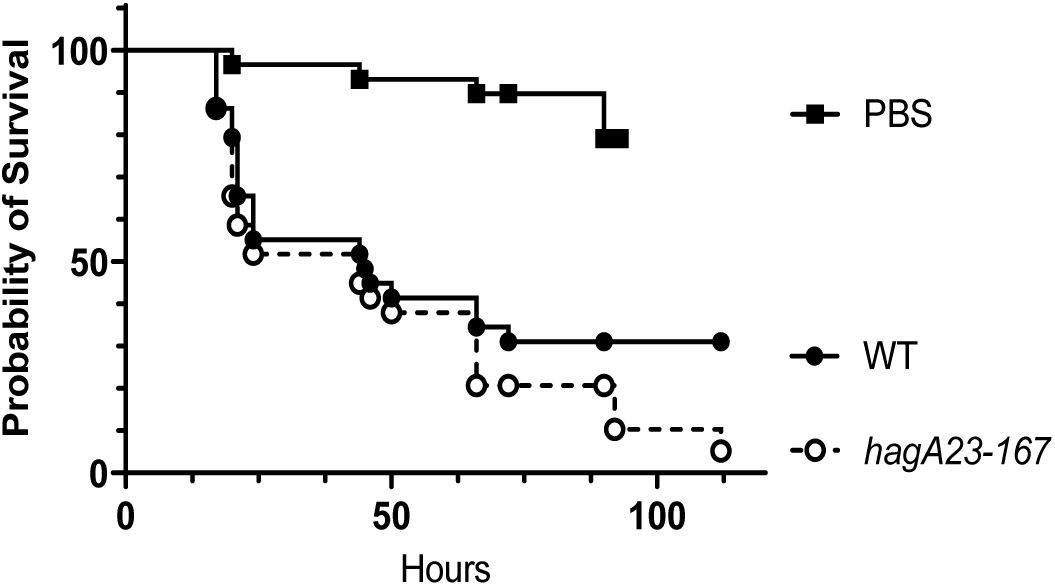
Survival of *G. mellonella* infected with *P. gingivalis*. 6^th^ instar larvae were injected with PBS, W50 (WT) or *hagA23/167* and incubated for up to 110 hours. Survival was determined at the times indicated. The data are from three independent experiments; N=29. The curves were compared by Mantel-Cox logrank test. WT differs from PBS (P<0.0001) but not from *hagA23-167* (P<0.25).

## Discussion

### Identification of the hagA mutant

*P. gingivalis* depends on heme for growth and virulence (15). Thus, there has been great interest in non-pigmented mutants, which are defective in heme uptake (16, 28, 30). As part of an effort to understand the activity of the AMP GL13K against *P. gingivalis*, we identified a novel non-pigmented mutant that showed a low MIC and two-fold extended MDK to DGL13K, consistent with tolerance to this peptide (**Table 4**) (57). In contrast, pigmented, WT cells were susceptible to DGL13K but resistant to LGL13K, as previously reported (43) (**Table 4**).

**Table 4.**
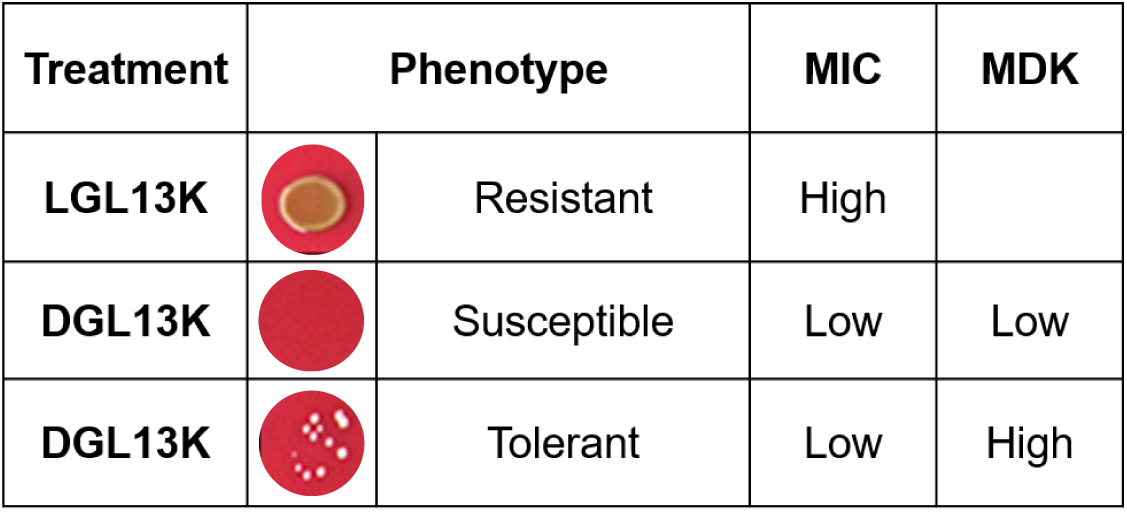
Activity of L- and D-enantiomers of GL13K against *P. gingivalis* W50. LGL13K exhibits a high MIC, suggesting that the bacteria are resistant to this peptide. DGL13K exhibits a low MIC and WT cells show a low MDK, suggesting that these bacteria are susceptible to DGL13K. A non-pigmented mutant survives about twice as long as WT cells, indicating that this mutant (*hagA23/167*) is tolerant to DGL13K.

| Treatment | Phenotype |  | MIC | MDK |
| --- | --- | --- | --- | --- |
| LGL13K |  | Resistant | High |  |
| DGL13K |  | Susceptible | Low | Low |
| DGL13K |  | Tolerant | Low | High |

Our finding that *hagA23/167* was defective in hemoglobin binding and hemagglutination (**Fig. 4**) but not in proteolysis (**Fig. 6**) is consistent with the ablation of HagA activity while preserving gingipain activity. It is not clear how the two SNPs in the *hagA* sequence affect protein expression. Although the predicted mRNA structure for WT *hagA* differed from *hagA23/167*, the mRNA levels appeared to be similar in WT and mutant strains, suggesting that differences in translation, rather than transcription, were responsible for the change in HagA activity. Indeed, changes in codon usage and mRNA structure, caused by the synonymous SNPs, could cause a change in translational efficiency (59).

The adhesion domains of HagA, RgpA and Kgp are involved in cell-cell interaction and biofilm formation (66). In *Bacillus subtilis*, biofilm formation has been linked to the utilization of specific Ser codons and reduced expression of the regulatory protein SinR (67). Since our mRNA modeling suggested that the synonymous SNP at a Ser codon was responsible for the observed reduction of HagA function, we propose that a serine sensor may also regulate biofilm formation in *P. gingivalis*. According to this model, depletion of HagA may allow bacteria to escape a biofilm that is under attack by antimicrobial peptides.

Black pigmentation and hemin binding previously have been linked to the presence of anionic LPS (A-LPS) in the outer membrane of *P. gingivalis* (25, 68). The structure of *P gingivalis* LPS also plays a role in bacterial resistance to the AMP polymyxin B (40, 69). Specifically, a lack of phosphate groups in lipid A reduces the electrostatic attraction between the cationic peptide and LPS (69). Indeed, a phosphatase mutant that is defective in dephosphorylation of lipid A exhibits increased sensitivity to polymyxin B (40). HagA contains a conserved LPS attachment site (70) and is anchored in the outer membrane by A-LPS (62, 71). Thus, it is possible that reduced levels of HagA at the cell surface in the *hagA23/167* variant also alter the cell surface complement of LPS and thereby affects bacterial sensitivity to AMPs and pigmentation.

It is not entirely clear how potential differences in cell surface charges, mediated by LPS, differentially affect the two GL13K enantiomers. However, it has been reported that the two peptides are not merely mirror images of one another (72). Thus, the four pK_a_ values of LGL13K are somewhat higher than the corresponding pK_a_ values of DGL13K and the latter peptide adopts its secondary structure more rapidly than the L-enantiomer (72). These structural and charge differences may affect the interaction of the peptides with the bacterial outer membrane.

Given the role of LPS and HagA in bacterial virulence (18, 73), it is surprising that the mutant bacteria were at least as virulent as the WT strain. It is possible that, despite the non-pigmented phenotype, these mutants compensate for the loss of a virulence factor by functional redundancy (74), *e.g.* as expressed by the gingipain/hemagglutinin protein family (20).

## Materials and Methods

### Bacterial strains

WT *Porphyromonas gingivalis* W50 was kindly provided by Dr. Massimo Costalonga, University of Minnesota School of Dentistry. The *P. gingivalis* mutants KgpΔIg-B (Δ*kgp*) (75) and Δ266 (Δ*porN*) (62) were created in a W83 background and kindly provided by Dr. Jan Potempa, University of Louisville School of Dentistry.

Unless stated otherwise for individual assays, *P. gingivalis* were routinely cultured in ‘Todd Hewitt Medium’ (THM) consisting of Todd Hewitt Broth (BD Bacto 249240) supplemented with 0.01% Hemin and 0.01% Menadione (both from Sigma-Aldrich, St. Louis, MO). Bacterial cultures were plated on THM-blood-agar consisting of 5% of sheep blood (Remel Lenexa, KS) in THM with 1.5% agar (Fisher Scientific). The bacteria were routinely cultured in an anaerobic chamber in an atmosphere consisting of 80% N_2_, 10% CO_2_ and 10% H_2_ at 37°C.

### Peptides

LGL13K and DGL13K were purchased from AappTec (Louisville, KY) or Bachem (Torrance, CA) at >95% purity. Peptide identity and purity were confirmed by the supplier by mass spectrometry and RP-HPLC, respectively. Polymyxin B was purchased from Sigma (St. Louis, MO).

Peptides were dissolved at 10 mg/ml in sterile 0.01% acetic acid and stored at 4°C until use, as described (53). Peptide batches were tested for antimicrobial activity by MIC assays against *P. aeruginosa* Xen41 (56).

### Timed-kill assay

*P.gingivalis* was cultured in THM for 72 h at 37°C followed by centrifugation at 6500 x g for 10 min at 4°C. The cell pellets were resuspended in 10 mM sodium phosphate, pH 7.4 at 10^7^ CFU/ml. The bacterial suspension (495 µl) was mixed with 5 µl of peptide stock solution (10 mg/ml) and incubated in an anaerobic chamber at 37°C. Five µl aliquots were removed from the reaction mixture at the times indicated and plated on blood agar until brown colonies emerged.

### Minimal Inhibitory Concentration (MIC)

*P. gingivalis* were cultured in THM until an approximate optical density at 600 nm (OD_600_) = 1 was reached. The bacteria were diluted in THM to 10^5^ CFU/ml. MICs were determined as previously described (56, 76). Briefly, 100 µl of bacterial stock (10^4^ CFU/well) were mixed with 20 μl of a 2-fold serial peptide dilution (concentration range: 1025 µg/ml – 1 μg/ml; control samples 0 µg/ml) in 0.01% acetic acid. The samples were incubated in 96-well polypropylene plates in an anaerobic chamber at 37°C for 3 days. The OD_630_ was determined in a Synergy HT plate reader (BioTek, Winooski, VT) and plotted against peptide concentration. The MIC was read as the lowest peptide concentration that prevented any bacterial growth.

### Genome sequencing

DNA samples for whole genome sequencing were prepared from colonies isolated from four independent experiments. Each experiment contained both untreated WT *P. gingivalis* W50 and W50 that had been treated with DGL13K for 60 min. Individual colonies were selected and subcultured in THM to stationary phase and genomic DNA prepared using the Masterpure Gram-positive DNA purification kit (Lucigen, Middleton, WI). The samples then were purified further using ZymoDNA Clean & Concentrator-5 kit (Zymo Research, Irvine, CA). DNA sample quality was verified by a fluorimetric Picogreen assay (>0.2 ng/µl).

Genomic library generation and MiSeq sequencing were performed by the University of Minnesota Genomics Center. Briefly, DNA libraries were prepared using the Nextera XT DNA sample preparation kit (Illumina, San Diego, CA, USA), according to the manufacturer’s specifications. Libraries were then sequenced using an Illumina MiSeq platform (2×300 bp) using Illumina’s SBS chemistry. The BBDuk tool for Geneious software version 168 9.1.8 was used to quality trim and filter Illumina adapters, artifacts, and PhiX from reads. Paired reads with quality scores averaging <6 before trimming or with a length <20 bp after trimming were discarded.

Remaining reads were mapped to the published W83 reference genome sequence (NCBI Reference Sequence: NC_002950.2) via BWA (0.7.17-r1188) to generate BAM files. Variant calling was done in parallel across all samples via Freebayes using a minimum variant frequency of 0.01 and minimum coverage of 34 reads. Polymorphism frequencies in each culture were determined and gated at >10% threshold.

### mRNA modeling

The mRNA secondary structure was modeled for a 493 nucleotide segment coding for the C-terminal Cleaved Adhesin Domain (PFAM: PF07675) in HagA. This domain was identified as the C-terminal orphan K3 domain by Dashper et al. (60) and contains the two synonymous SNPs identified in this study. Centroid structures (77) were calculated using the default parameters in the Vienna RNA web suite (61, 78), which is available from http://rna.tbi.univie.ac.at/cgi-bin/RNAWebSuite/RNAfold.cgi

### Reverse transcription-polymerase chain reaction (RT-PCR)

*P. gingivalis* were cultured for 48 h, as described above. Bacteria were centrifuged for 10 min at 10,000 x g (4°C) and the cells resuspended in 0.5 ml of TRIzol (Thermo Fisher Scientific). Direct-zol, RNA Miniprep Plus and RNA clean-up Quick-RNA Miniprep Kits (Zymo Research) were used to isolate and purify mRNA. The mRNA was converted to cDNA using the Invitrogen PhotoScript II First-Strand Synthesis System (New England Biolabs, Ipswich, MA) according to the manufacturer’s instructions.

PCR reactions were conducted in a PCR Mastercycler (Eppendorf North America, Hauppauge, NY). A 20 μl reaction containing 1.0 μl of cDNA, 10µM of each primer and 10 μl of OneTaq 2X Master Mix with Standard Buffer (New England Biolabs) was used for double-strand DNA synthesis. The RT-PCR reactions were carried out following the recommended thermal profile: 94°C for 5 min, followed by 35 cycles of 94°C for 45 s, 58°C for 45 s and 72°C for 2 min. The specificity of the amplicons was determined by electrophoresis of the PCR products on 1.5% agarose gels (BioExpress, Chicago, IL).

Primers for *P. gingivalis* 16S rRNA: 5’-TGTAGATGACTGATGGTGAAAACC-3’and 5’-ACGTCATCCCCACCTTCCTC-3’; Primers for hagA: 5’-GGAGGCTCACGATGTATGGG-3’ and 5’-ATCGGCATTGACCGGAACTT-3’.

### Hemoglobin binding

*P. gingivalis* cultures were centrifuged at 10,000 x g for 10 min and the cell pellets resuspended in PBS to approximately OD_600_ = 1. Human red blood cells (1ml) were centrifuged 10 min at 10,000 x g. The pellets were resuspended in an equal volume of dH_2_O and then frozen at -20°C. The cells were thawed at room temperature and centrifuged as before. The supernatant (RBC lysate) was used for hemoglobin binding. Aliquots of bacterial suspension (750 µl) were mixed with 250 µl of RBC lysate. One set of samples was immediately centrifuged for 10 min at 10,000 x g (0 h incubation). A second set of samples was incubated for 1 h at 37°C and then centrifuged (1 h incubation). The supernatants were collected and the OD_450_ read in a plate reader. Hemoglobin binding was calculated as (OD_450_@0h – OD_450_@1h) / OD_450_@0h

### Hemagglutination

This assay was based on published protocols (26, 79). Human red blood cells were washed twice in PBS and resuspended in 10x their original volume. Cultures of *P. gingivalis* (2 days) were centrifuged at 10,000 x g for 10 min and the pellets resuspended in PBS to an approximate OD_600_ = 1. Aliquots of the bacterial culture (80 µl) were diluted 2-fold in 80 µl PBS in round bottom polyvinyl chloride 96-well plates. Diluted red blood cells (80 µl) were added to each well and the plates were incubated for 3-4 h at 37°C with rocking on a Bellydancer laboratory shaker (Stovall Life Science, Greensboro, NC). Hemagglutination was determined as the highest dilution of the bacterial culture that caused ‘matting’ of the red blood cells.

### Whole-cell ELISA

*P. gingivalis* was cultured on hydroxyapatite-coated pegs placed in a 96-well plate (MBEC biofilm incubator; Innovotech, Edmonton, AB, Canada) for 48h. The pegs were rinsed in 300 µl/well of PBS and transferred to a polypropylene 96-well plate containing 200 µl/well of PBS with 0.05% Tween 20 and 0.5% non-fat powdered milk and the monoclonal antibody 61BG1.3 diluted 1:10,000. This antibody was developed by R. Gmuer, Institute of Oral Biology, ZZMK, University of Zürich, Switzerland. It was obtained from the Developmental Studies Hybridoma Bank, which was created by the NICHD of the NIH and maintained at the University of Iowa, Department of Biology, Iowa City, IA.

The plates were incubated with the antibody for 2.5 h at 30°C with gentle shaking and then rinsed twice in PBS followed by a 20 min wash in 300 µl/well of PBS. Each well was then incubated for 90 min at 30°C with 200 µl PBS containing 0.05% Tween 20, 0.5% dry milk and horseradish peroxidase-conjugated rabbit anti-mouse IgG diluted 1:1,000. The pegs were washed in PBS as before and incubated with horseradish peroxidase substrate in a 96-well white wall polystyrene plate. Luminescence was measured in a Biotek Synergy HT platereader after 15 min incubation with ECL Western Blotting substrate (Thermo Pierce, Rockford, IL). For each strain, the average luminescence readings in four wells, which had been incubated without primary antibody, were subtracted from each antibody reading.

### Hemoglobinase Assay

Stationary phase cultures of *P. gingivalis* were centrifuged for 10 min at 10,000 x g. The supernatants (180 µl) were mixed with 20 µl of 5 mg/ml hemoglobin in PBS. The OD_405_ was read at T = 0 h and then the plates were incubated for 24 h at 37°C and OD_405_ determined. The net OD_405_ was determined by subtracting background values obtained in the absence of hemoglobin at each time point. Relative proteolysis was calculated as netOD_405_@0h – netOD_405_@24h.

### *In vivo* virulence

Stationary phase cultures of *P. gingivalis* were centrifuged for 10 min at 10,000 x g and the cells were resuspended in PBS to approximately 10^6^ CFU/ml. Sixth instar larvae of *Galleria mellonella* (greater wax moth) were purchased locally and kept in sawdust at 4°C until use (typically 1-3 days). Groups of 9-10 larvae (270 ± 4 mg each) were injected with 10 µl (10^4^ CFU) of *P. gingivalis* and incubated in a polystyrene Petri dish with oatmeal at 37°C in the dark. Dead larvae were identified by melanization and absence of movement when prodded (56, 65). Surviving larvae were counted at the times indicated.

### Data availability

All data are contained within the manuscript

## Acknowledgements

Dr. Helmut Hirt, University of Minnesota is acknowledged for the initial discovery of non-pigmented colonies upon DGL13K treatment. We thank Dr. Donald Demuth, University of Louisville for helpful suggestions and Dr. Pamela Hanic-Joyce, Concordia University, Montreal for helpful suggestions and careful reading of the manuscript. Dr. Massimo Costalonga, University of Minnesota School of Dentistry is thanked for providing *P. gingivalis* W50 and Dr. Jan Potempa, University of Louisville is thanked for making gingipain mutants available for this work.

## Funding

This work was supported by an Academic Health Center Seed Grant from the University of Minnesota and research funds from the University of Minnesota School of Dentistry.

### Conflict of Interest

The authors declare no conflicts of interest in regards to this manuscript

**Footnotes**

## Abbreviations

AMP: antimicrobial peptide
CFU: colony-forming units
LPS: lipopolysaccharide
MDK: minimum duration for killing
MIC: minimum inhibitory concentration
OD: optical density
THM: Todd-Hewitt medium

